# Activity of Telavancin against *S. aureus* isolated from cystic fibrosis patients including those with decreased susceptibility to Ceftaroline

**DOI:** 10.1101/319210

**Authors:** Melanie Roch, Maria Celeste Varela, Agustina Taglialegna, Warren E. Rose, Adriana E. Rosato

**Author notes:** Corresponding author: Houston Methodist Research Institute, 6670 Bertner Ave., Room R6-113, Houston, TX 77030.

## Abstract

Methicillin-resistant *Staphylococcus aureus* (MRSA) acquisition in cystic fibrosis (CF) patients confers a worse clinical outcome with increased rate of declined lung function. Telavancin, an approved lipoglycopeptide used to treat infections due to *S. aureus* has a dual mode of action causing inhibition of the peptidoglycan synthesis and membrane depolarization. CF-associated MRSA infections remain an important problem with no foreseeable decline in prevalence rates. Although telavancin is currently in clinical use for complicated skin infections and hospital-acquired pneumonia, the activity against CF-associated *S. aureus* infections has not been investigated. In this work, we studied the activity of telavancin against CF *S. aureus* strains collected from diverse geographical CF centers in the USA. We found that telavancin-MIC90 was 0.06 μg/ml, 8-fold lower than ceftaroline or daptomycin and 25-fold lower than linezolid and vancomycin. We demonstrate that telavancin at serum-free concentrations has rapid bactericidal activity with a decrease of more than 3 log10 CFU/ml during the first 4 to 6 hours of treatment, performing better in this assay than vancomycin and ceftaroline, including *S. aureus* resistant to ceftaroline.

Telavancin resistance was infrequent (0.3%), although we found that it can occur *in-vitro* in both CF-and non-CF *S. aureus* strains by progressive passages with sub-inhibitory concentrations. Genetic analysis of telavancin *in-vitro* mutants showed gene polymorphisms in cell wall and virulence genes, and increased survival in a *Galleria mellonella* infection model. Thus, we conclude that telavancin represents a promising therapeutic option for CF infections with potent *in-vitro* activity and low resistance potential.

## INTRODUCTION

Methicillin-resistant *Staphylococcus aureus* (MRSA) is an important infectious human pathogen responsible for diseases ranging from skin and soft tissue infections to life-threatening endocarditis, both in hospitals- (HA) and community- (CA) acquired settings. β-lactam resistance in MRSA involves the acquisition of penicillin-binding protein (PBP) 2a, which has low affinity for β-lactams and can mediate cell wall assembly when the normal staphylococcal PBPs (PBP1 to 4) are inactivated by these agents (1). *S. aureus* is one of the earliest and more prevalent pathogens colonizing respiratory tracts and then causing infection in people with cystic fibrosis (CF). This severe, autosomal recessive disease affects several organs, notably the lungs, predisposing these patients to reduced respiratory function and infections with potentially severe consequences. According to recent data from the CF Foundation Patient Registry, the prevalence in USA of MSSA is around 70%, while MRSA is 26%. These values compare to 13% in Europe, 6% in Canada and 3% in Australia (2, 3). Emerging research has demonstrated that MRSA infections have a significant clinical impact on individuals with underlying chronic diseases such as CF, where antibiotic pressure and metabolic adaptions may favor the ability of *S. aureus* to establish long persistence and resistance (4). Additional mechanisms reported in CF lung and other chronic MRSA infections reduce antibiotic activity including small colonies variant adaptation, biofilm formation and growth under anaerobic conditions, which are associated with higher rates of antimicrobial treatment failure (5–8). Moreover, in CF, chronic pulmonary infections with MRSA and their exacerbations were shown to be associated with decline in lung function and a worse clinical outcome. (9) In this context, additional data regarding the antibiotic susceptibility of strains from CF-patients are urgently needed to enhance treatment options against multi-drug resistance and to try to eradicate MRSA from their lungs.

Telavancin (TLV) is a lipoglycopeptide antibiotic approved by the FDA in 2009 for the treatment of complicated skin and skin structure infections and in 2013 for nosocomial pneumonia (HAP/VAP) (10) suspected to be caused by MRSA. TLV was developed from the parent molecule vancomycin (11) and its bactericidal action involves membrane depolarization and the inhibition of the peptidoglycan (PG) synthesis (12, 13), at the late stage PG precursors, including lipid II; however, the precise mode of action of TLV on Gram positive membranes has not yet been determined (6–8).

*In vitro* studies have demonstrated TLV activity against MRSA, including vancomycin-intermediate *S. aureus* (VISA) strains (10, 14). However, no clinical data is yet available about TLV activity against MRSA strains isolated from patients with chronic diseases such as cystic fibrosis. These strains are well known to have altered metabolism and possess multi-drug resistance due to their chronic habitation and prolonged, repeated exposures to antibiotic treatments in the CF lung environment (15).

We hypothesized that TLV may represent a valid option for the treatment of CF-MRSA and MSSA infections. Therefore, the purpose of this study was to characterize by *in-vitro* and *in-vivo* approaches the antimicrobial activity of TLV in *S. aureus CF* chronic infections strains, particularly, infections by MRSA, strains isolated from diverse CF centers in the United States. Lastly, we aimed to understand TLV resistance selection within a *S. aureus* population derived from a CF-background.

## MATERIAL AND METHODS

### Clinical CF strains

CF-strains were isolated from patients’ sputum cultures. A collection of strains comprising either wild type or small colony variant (SCV) phenotypes was obtained from three academic medical institutions with large CF populations: the Center of Global Infectious Diseases (Seattle, WA), UW Health (Madison, WI) and the Houston Methodist Hospital (Houston, TX).

### Susceptibility testing

TLV, ceftaroline (CPT), daptomycin (DAP), linezolid (LZD) and vancomycin (VAN) susceptibilities were determined by E-test (BioMerieux). TLV minimal inhibitory concentrations (MICs) were also determined by microdilution method in Mueller Hinton Broth II cation adjusted (MHBII) broth supplemented with polysorbate 80 (0.002%) following the CLSI guidelines (16).

### Time-Kill Analyses

Analyses were performed on 4 representative CF-strains following CLSI guidelines using human free drug maximal concentrations for TLV (8 mg/L for 750 mg dose, VAN (16 mg/L for a 1000 mg dose), LZD (10.4 mg/L for a 600 mg dose) and CPT (16 mg/L for a 600 mg dose) in 24 well–microplates (17–20). Colony forming units (CFU) were counted by plating a sample on Tryptic Soy Agar (TSA) at 0, 2, 4, 6, 8 and 24 hours. Results were expressed in CFU/ml *vs.* time.

### TLV *in-vitro* mutant selection

*In-vitro* selection was attempted in a representative number of CF-strains. *S. aureus* ATCC 25923 was included as a non CF-reference strain. Strains and controls were exposed to sub-inhibitory concentration of TLV in MH during 40 days by progressive passages to recover non-susceptible TLV strains.

### Assessment of virulence between TLV susceptible and resistant mutants in a Wax worm model of infection

Groups of *Galleria mellonella* larvae (10/group) were inoculated with 10 μl of a bacterial suspension of strains AMT 0114-48, AMT 0114-48 TLV –R, and ATCC 25923, ATCC 25923 TLV-R strains, containing 1.5 x 10^6^ CFU/ml, as previously described (21). Inoculum was administered directly to the larval hemocoel through the last left pro-leg as previously described (17, 22). Every trial included a group of 10 untreated larvae as uninfected control group and 10 larvae injected with PBS as a method control. Experiments were performed in at least three independent trials. Injected insects were monitored over seven days at 37°C. By the day seven, pupa formation was recorded in survived larvae.

### Genomic Characterization and Whole Genome Sequencing

DNA from the different strains were prepared using the DNeasy Blood & Tissue Kit (Qiagen). Libraries were prepared from purified DNA using Nextera XT DNA Library Preparation Kit (Illumina) and sequenced with HiSeq 2000 instruments at the Epigenetics and Genomic laboratory at Weill Cornell University, NY. Genomes were assembled and SNPs were identified and compared to *S. aureus* N315 (GenBank BA000018) using the Lasergene 14 Suite. Reads were aligned against N315 (PATRIC ID: 158879.11) and analyzed using the PATRIC variation service 24, which uses BWA-mem as the read aligner (https://arxiv.org/abs/1303.3997) and FreeBayes as the SNP caller (https://arxiv.org/abs/1207.3907).

## RESULTS

### Susceptibility of Cystic Fibrosis-derived MRSA/MSSA strains to Telavancin

The collection of CF-*S. aureus* strains used in the present study were either wild type or small colony variants (SCV) phenotype identified at culture collection, and were obtained from three different CF centers. We screened a total of 333 strains, those collected at Houston Methodist Research Institute, 103 in total, were distributed as 37 MRSA and 66 MSSA; while strains originating from the UW Health comprised 72 MRSA and 10 MSSA strains. The ones obtained from the Center of Global Infectious Diseases at Seattle (WA) comprised 148 *S. aureus* total strains with 44 MRSA and 104 MSSA. As shown in Table 1, TLV displayed activity against all the 333 CF strains derived from 3 different CF centers across the USA; the majority of these *S. aureus* were isolated from adult patients, with only 41 strains found in children. Furthermore, we tested the activity of TLV against 23 MRSA strains; 20 displaying intermediate resistance to CPT (CPT-IR, MIC: 1.5-2 μg/ml) and three with high level of resistance (CPT-HR; MIC: 32 μg/ml) strains.

**Table 1.**
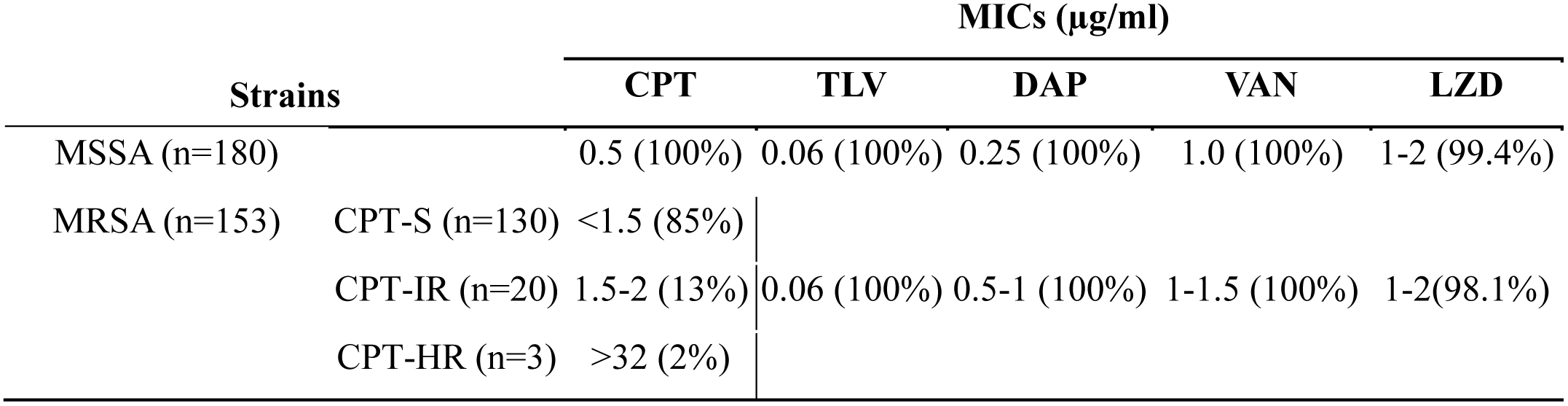
Telavancin MICs 90 performed in 333 CF-*S. aureus* strains determined by microdilution method in Mueller-Hinton broth supplemented with polysorbate 80 (0.002%) and in comparison to DAP, VAN, LZD and CPT agents following the CLSI guidelines.

| Strains |  | MICs (µg/ml) |  |  |  |  |
| --- | --- | --- | --- | --- | --- | --- |
|  |  | CPT | TLV | DAP | VAN | LZD |
| MSSA (n=180) |  | 0.5 (100%) | 0.06 (100%) | 0.25 (100%) | 1.0 (100%) | 1-2 (99.4%) |
| MRSA (n=153) | CPT-S (n=130) | <1.5 (85%) |  |  |  |  |
|  | CPT-IR (n=20) | 1.5-2 (13%) | 0.06 (100%) | 0.5-1 (100%) | 1-1.5 (100%) | 1-2(98.1%) |
|  | CPT-HR (n=3) | >32 (2%) |  |  |  |  |

CPT is a new β-lactam antibiotic that specifically targets PBP2a in MRSA. Although high level of resistance to CPT remains rare, intermediate resistance is more prevalent in patients with chronic infections. Among all strains, TLV-MIC_90_ was 0.06 mg/L, i.e. 8-fold lower than DAP and CPT, and 25-fold lower than LZD and VAN. In the strains with reduced CPT susceptibility, the TLV MIC_90_ was 0.06 μg/ml for both CPT-IR TMH-5006 and CPT-HR TMH-5007 strains, showing the absence of cross-resistance between both antibiotics (Table 1). Of note, although DAP showed *in-vitro* activity against CF-*S. aureus* strains, in the presence of 1% surfactant resulted in a 8 to 32 fold increase in DAP MIC (up to 8 mg/L), while the MICs for the other antibiotic, including TLV, remain unchanged (data not shown). These data support previously documented data showing inactivation of daptomycin in the lung, and provide evidence that other MRSA treatments options retain potency.

Of note, we found in AMT-0067-21 strain the presence of internal colonies in TLV E-test for strain and displaying (MIC: 0.19 μg/ml; homogeneous up to 0.047 μg/ml) and a broth microdilution MIC at 0.25 μg/L. Interestingly, the tested MRSA and MSSA strains were not associated to VISA or DAP non susceptible phenotype. These data altogether provide evidences that TLV retain potency against S. aureus CF strains.

### TLV shows bactericidal activity against CF-MRSA/MSSA isolates

The *in-vitro* effectiveness of TLV was also evaluated by time-kill experiments compared to DAP,VAN and CPT. The assay was performed in Muller-Hinton broth supplemented with 0.002% polysorbate-80 for TLV and calcium 50 μg/ml for DAP, against a representative number of CF-strains including a high CPT-R (TMH-5007) strain. (18–20). TLV showed activity against all the tested strains including the CPT-HR strain TMH 5007 in concordance with TLV MIC values. Moreover, TLV displayed rapid bactericidal activity with a decrease of more than 3 log10 CFU during the first 4 hours of growth (Figure 1). Similar activity was observed in the CPT-HR strain confirming the absence of cross-resistance or reduced activity. TLV activity profile at free serum concentration 8 mg/L performed better than VAN (16 mg/L), LZD (10.4 mg/L) and CPT (16 mg/L) (Figure 1). Together these data support the potential therapeutic application of TLV for the treatment of *S. aureus* CF infections.

**Figure 1.**
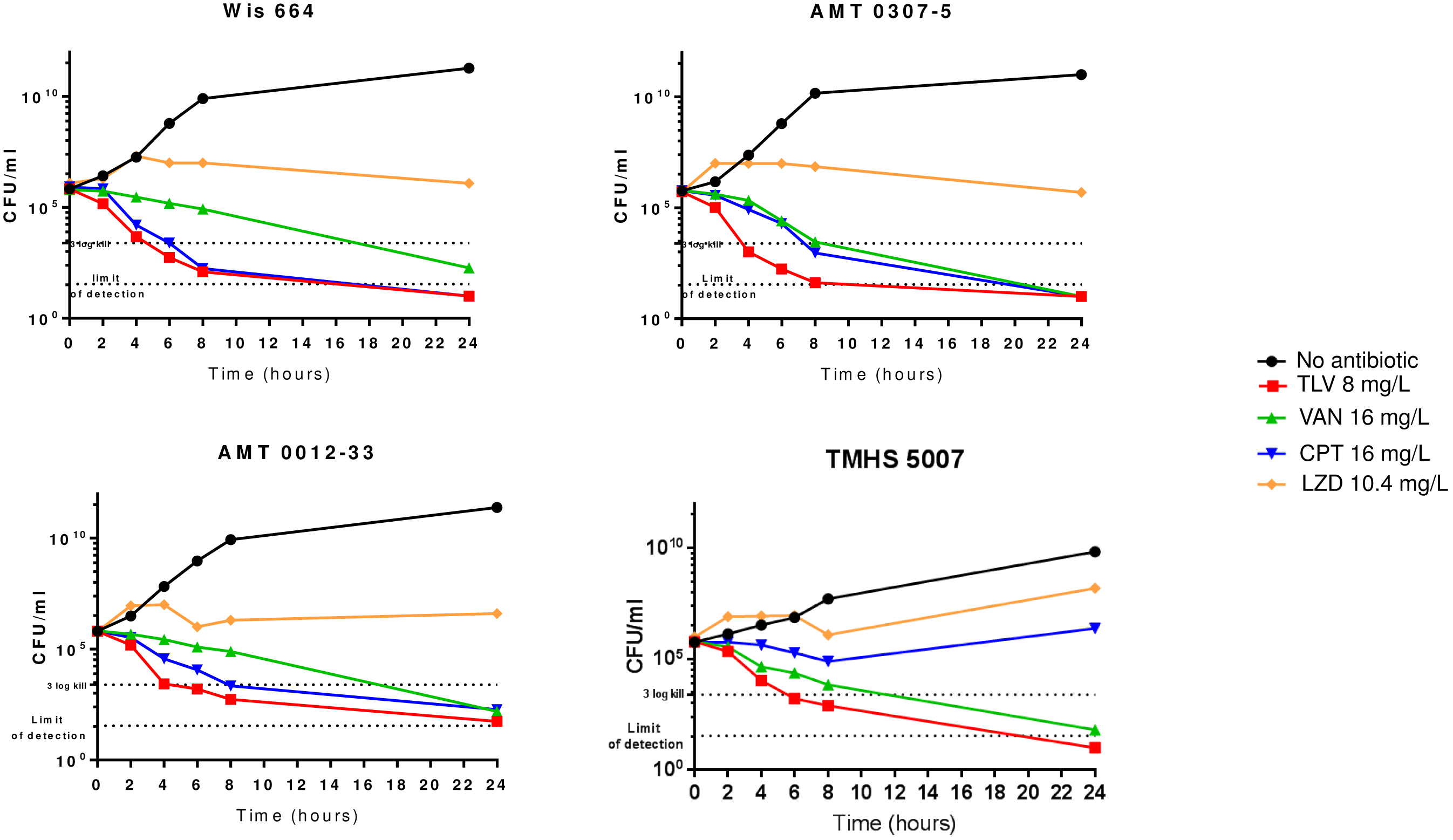
Time-kill curves of CF-strains AMT 0307-5 (MSSA), AMT0012-33; WIS 664 (MRSA) and TMHS-5007 (MRSA CPT-R) strains and using human peak free serum drug concentrations: TLV=8mg/L, VAN=16mg/L, LZD=10.4mg/L and CPT=16mg/L. The limit of detection of that assay was 10.

### *In-vitro* selection of TLV resistant mutants

To determine the fate of mutation selection that can be projected by the potential prolonged use of TLV in CF patients we investigated the ease of *in-vitro* selection from three clinical CF representative strains: AMT 0114-8, WIS 664 and TMH5007 (CPT-HR) strain. *S. aureus* ATCC 25923 was included as a non-CF strain control. These strains were serial passaged at sub-inhibitory concentrations of TLV starting at 0.03 μg/ml and escalating up to 3μg during 40 days. After 15 days, the strains showed an increase in TLV MIC from 0.06 μg/ml up to 0.25 -1 μg/ml, followed by a progressive increase up to 3 μg/ml after 40 days of exposure (Table 2). The enhanced MICs of the mutants were stable and unaltered after ten passages in the absence of TLV. As shown in Table 2, there was 3 to 4 fold increase in VAN MIC going from 1.5 to 6 μg/ml and an 8 to 10 fold increase in DAP MICs suggesting potential cross-resistance between TLV, VAN and DAP antibiotics. Moreover, the *in-vitro* TLV mutants grew at slower rate than the parent strains and were defective in growth requiring 48h to obtain normal size colonies on TSA blood agar plates (data not shown). Important observations are taken from these results: (i)-the ease of mutant selection observed in *S. aureus* ATCC 25913 control strains lead us to conclude that TLV mutant resistance is independent of CF background strains, and (ii) the likelihood of TLV strains with increased TLV –MICs (0.19 μg/ml) as observed for only one *in-vivo* strain MT-0067-21 seems rare considering the fact that this strain represent only the 0.3% of the total 333 tested strains.

**Table 2.** MICs of TLV, VAN and DAP of parents and *in-vitro* TLV mutants obtained by serial passage with sub-inhibitory concentrations of the antibiotic (TLV) during 40 days. As shown, a 3-4 fold increase in VAN MIC (from 1.5 to 6 μg/ml) and an 8-10 fold increase in DAP MICs (0.25 to 8 μg/ml) were determined, suggesting potential cross-resistance between TLV, VAN and DAP antibiotics.

| STRAINS | DAY 0 |  |  | DAY 40 |  |  |
| --- | --- | --- | --- | --- | --- | --- |
|  | TLV | VAN | DAP | TLV | VAN | DAP |
| ATCC 25913 | 0.064 | 1.5 | 0.25 | 3 | 4 | 8 |
| TMH 5007 | 0.047 | 2 | 0.75 | 1.5 | 6 | 8 |
| AMT 0114-48 | 0.047 | 2 | 0.75 | 2 | 6 | 6 |
| WIS 664 | 0.032 | 2 | 1 | 1.5 | 3 | 6 |

### Genes associated to *in vivo* and *in vitro* TLV resistance

To investigate the genetic mutations associated with the *in-vivo* (MRSA - MT-0067-21) or *in-vitro* TLV resistant mutants, all the strains were full genome sequenced and compared with the reference strain *S. aureus* N315 (PATRIC ID: 158879.11). Mutated genes were categorized by function to identify themes of bacterial physiology that may contribute to reduced susceptibility to TLV (Table 3). The most common non-synonymous SNP found by comparing each strain with their counterpart parental strain cell wall associated genes correspond to *sdr*CDE (cell wall associated genes), *tcaA* (trans-membrane protein associated with teicoplanin resistance), *dltD* (D-alanine transfer protein). In addition, we found additional non-synonymous SNPs in various cell wall associated genes: *spa*, *clfA, clfB, sdrE and srdD* (Table 3). Moreover, additional SNPs were found in the *in-vivo* derived strain MRSA MT-0067-21) displaying reduced TLV activity, notably *murE* (stop codon at position Q), *dltA, vraG* (bacitracin export permease). Gene deletion occurred on *isdB* (cell surface receptor for hemoglobin), *mutL* (DNA mismatch protein), *fnB* (fibrinogen binding protein) and in the *YycFG* regulon (two component regulatory system) (Table 3). These results may suggest that decrease susceptibility to TLV is mainly associated to changes in cell wall related genes.

**Table 3:**
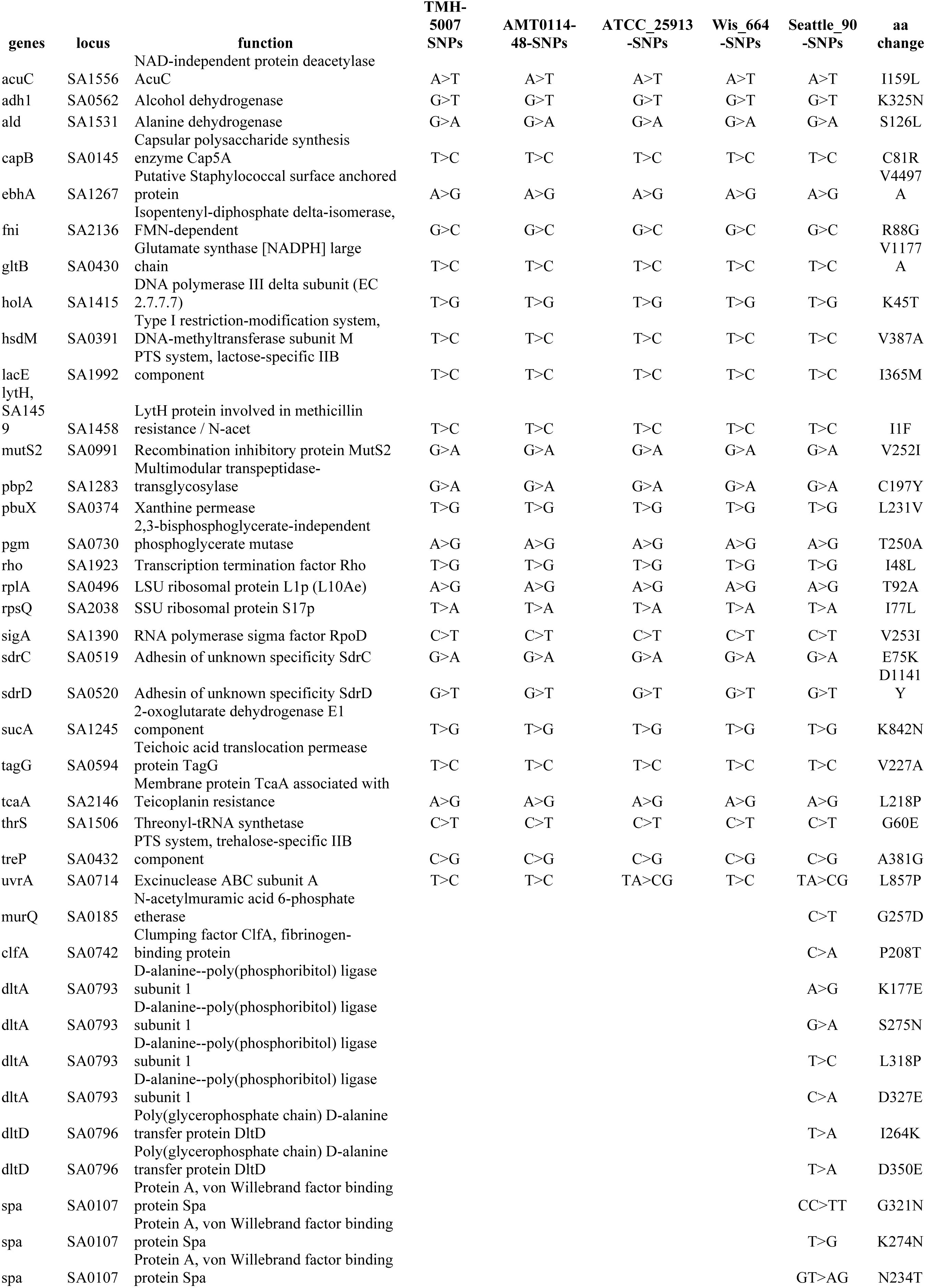
Most relevant mutations identified in telavancin resistant mutants

### *In-vitro* TLV mutants are associated with reduced virulence

In order to determine whether TLV resistance acquired *in-vitro* may impact virulence traits in *S. aureus* we used a *Galleria mellonella* as an *in-vivo* model that possess an immune system with reasonable homology to that of vertebrates, and numerous enzymatic cascades akin to complement fixation and blood coagulation occur in the hemolymph, resulting in hemolymph clotting and melanin production as key defense mechanisms against invading microbes. These tissue types are similar to those encountered by *S. aureus* during invasive infections in humans (22).

Groups of larvae (10/group) were inoculated with a bacterial suspension containing parent strains AMT 0114-48, WIS 664 and a control ATCC 25913 and their corresponding TLV *in-vitro* derivative mutants AMT 0114-48 TLV-R, WIS 664-TLV-R and ATCC 25913-TLV-R (10^6^ bacteria/worm) as previously described (22). An uninfected control group received PBS treatment to control for multiple injections. Worms were daily monitored and recorded for any deaths during 10 days. Worms injected with PBS showed 100/90% survival at day 8 (Figure 2), but groups of parent strains (e.g. AMT 0114-8) displayed low survival rates (≤50–0%, day 6; Figure 2). By contrast, groups of worms infected with TLV-R strains (e.g. AMT 0114-48-TLV-R) resulted in survival rates of 90% at day 6 followed by 60% survival at day 8. Similar trend was observed with the ATCC 25923 strain although the survival rate was higher (40%) than parent *S. aureus* CF strain (0-20%) while TLV-R strains were comparable in survival. These results may suggest that TLV-R is associated with changes in virulence fitness.

**Figure 2.**
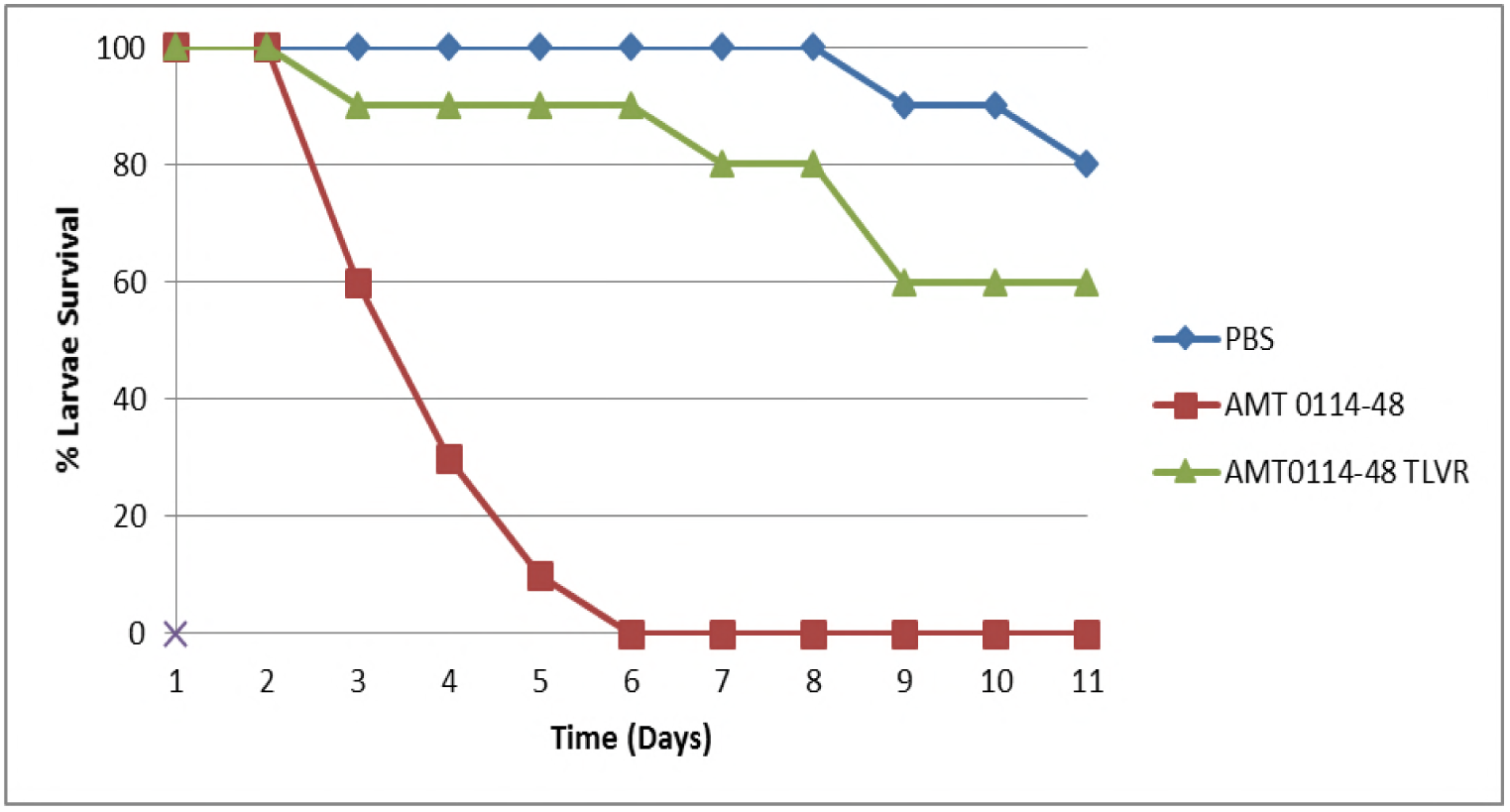
Comparison of virulence of *S. aureus* strains and their isogenic TLV mutants in *G. mellonella* model of infection .Parent strains ATCC 25923 and AMT 0114-18 are indicated in red line, mutants ATCC 25923 –TLVR and AMT0114-48 TLVR, indicated in green, infected at doses 10^6^ bacteria/worm and PBPS uninfected worms ( blue). Survival was monitored during 10 days.

## DISCUSSION

Infection with *S. aureus* remains an important concern for CF patients with consistently high prevalence in this population. Chronic MRSA infections associated with worse outcomes and treatment eradication is a continual clinical challenge. For MRSA pneumonia the most widely recommended antibiotics are vancomycin and linezolid, with TLV also approved for this infection type. Although the efficacy of TLV has been proven in VAP and HAP, less is known about the activity and potential efficacy in CF-patients with *S. aureus* pneumonia. In this sense, our study was performed retrospectively to assess the activity of TLV in *S. aureus* isolated from three different CF patient centers. The majority of the samples were collected from sputum and from both adults and children during the period from 2015-2017. We found that TLV was active against the majority of tested strains with exception of one that showed a slightly decreased activity (MICs; 0.19 μg/ml) above the TLV breakpoint.

There are no guidelines or recommendations on the choice of antibiotics for CF pulmonary exacerbations with MRSA, resulting in variable use of active antibiotics between centers. The most frequently used therapies in current practice are TMP-SMX (30%) linezolid ( 27%) and vancomycin ( 30%), with LDZ and VAN the most frequently used among inpatients (15, Zobell, 2015 #1020)The pharmacokinetic data available in healthy subjects on TLV intrapulmonary diffusion showed a good penetration of TLV into epithelial lining fluid and an extensive penetration into alveolar macrophages (23). A clinical trial is currently being conducted (https://clinicaltrials.gov/ct2/show/NCT03172793) to evaluate the pharmacokinetic profile of this drug in CF-patients who usually need dose adjustment due to an increase volume of distribution and clearance. TLV is not impacted by the pulmonary surfactant, unlike daptomycin, making it suitable for the treatment of CF associated lung infections. These observations are consistent with our results. In fact, we demonstrated that TLV has bactericidal activity against the *S. aureus* strains tested including those displaying reduced CPT and LZD activity (e.g. TMHS 5007) which might be a significant advantage, compared to the drug currently used, to try to eradicate those strains and prevent future exacerbations.

In absence of suitable alternative, TLV has been already used in some CF patients with successful outcomes (24), supporting its potential role in the management CF-MRSA infections. We are unaware of the development of TLV resistance in clinical settings. While direct resistance may be infrequent, modest increases in MICs may be seen in some isolates as the one described here (TLV MIC 0.19 μg/ml) and in some strains with VAN and DAP decreased susceptibility. In this context, it was advantageous to gain an understanding of the ease of TLV resistance when CF strains are extensively exposed to sub-therapeutic TLV *in-vitro* exposure. After 15 days, the strains showed an increase in TLV MIC from 0.06 μg/ml up to 0.25 -1 μg/ml, which is above the susceptible breakpoint of 0.12 μg/mL for *S. aureus*, followed by a progressive increase up to 3 μg/ml after 40 days of exposure. These observations may imply that resistance should be monitored in patient receiving repeated and/or prolonged treatment by TLV. In a previous study performed by (25) one stable mutant was obtained after 43 days in multistep resistance selection studies from one of the ten MRSA strains, correlating with low mutation frequency. Interestingly in our study, we were able to demonstrate that the TLV resistance selection was independent of the CF –*S. aureus* background strains considering the fact that we were able to obtain a TLV-R mutant from an ATCC 25923. We determined next the main genetic changes associated to TLV-R with full genome sequencing of a representative number of *in-vitro* obtained TLV-R strains along with the *in-vivo* derived CF-TLV-R that showed a modest increase of MIC (0.19μg/ml). We found that TLV-R harbored common mutations in genes associated to cell wall (e.g. *murE, tagB*) and cell wall surface (*spa, clfB, sdrE*) genes. In the TLV obtained in previous studies by Song Y et al (26)most of the differential expressed genes were also associated to changes in cell envelope. These findings suggest that although TLV is an agent with dual mechanism (cell membrane/cell wall) the compensatory preferential mutational mechanism seems to be in higher degree linked to cell wall (CW). These cell wall mutations may also function in a dual manner to reduce TLVs cell membrane mechanism potency. This is evidenced by prior studies correlating mutations and reduced VAN susceptibility as a result of VAN treatment (VISA strains) with collateral reduced DAP activity. Similarly, in our study the derived TLV-R mutants showed cross-resistance with reduced susceptibility to both VAN (MIC 3-6 μg/ml) and DAP (MIC 6-8 μg/ml).

Another set of genes in were we found non-synonymous SNPs were related to virulence (e.g. *sdrE*, *spa, clfB)*. These changes appeared to suggest that TLV-R affects *S. aureus* virulence fitness as evidenced in groups of worms infected with TLV-R strains (e.g. AMT 0114-48-TLVR) which resulted in increased survival rates (90%) compared to parent strains that manifested low survival rate. In works performed by (25) was shown a decrease expression of various virulence factors, however their functional role was not demonstrated.

In conclusion, the present data suggest that TLV is active against CF-MRSA strains independently of associated CPT resistant mechanisms and may constitute a new option for the treatment of CF-MRSA infections

## ACKNOWLEDGEMENTS

We acknowledge Dr Maria P. Martinez and Mrs. Maryam Fatouraei for their technical support. Drs. Rafael Hernandez, Luke Hoffman (Seattle, WA) for providing strains through their CF center strain repository and Dr. Wesley Long from Houston Methodist for providing CPT-HR resistant and CF strains. We also acknowledge the Epigenomics Core Facility at Weill Cornell Medicine (New York, NY, United States) for their resources and assistance with whole genome sequencing experiments.

